# A multiplexed time-resolved fluorescence resonance energy transfer ultrahigh-throughput screening assay for targeting SMAD4-SMAD3-DNA complex

**DOI:** 10.1101/2023.07.15.549169

**Authors:** Wukun Ouyang, Qiankun Niu, Min Qui, Haian Fu, Yuhong Du, Xiulei Mo

## Abstract

The signaling pathway of transforming growth factor-beta (TGFβ) plays crucial roles in the establishment of an immunosuppressive tumor microenvironment, making anti-TGFβ agents a significant area of interest in cancer immunotherapy. However, the clinical translation of current anti-TGFβ agents that target upstream cytokines and receptors remains challenging. Therefore, the development of small molecule inhibitors specifically targeting SMAD4, the downstream master regulator of TGFβ pathway, would offer an alternative approach with significant therapeutic potential for anti-TGF-β signaling. In this study, we present the development of a cell lysate-based multiplexed time-resolved fluorescence resonance energy transfer (TR-FRET) assay in an ultrahigh-throughput screening (uHTS) 1536-well plate format. This assay enables simultaneous monitoring of the protein-protein interaction (PPI) between SMAD4 and SMAD3, as well as the protein-DNA interaction (PDI) between SMADs and their consensus DNA binding motif. The multiplexed TR-FRET assay exhibits high sensitivity, allowing the dynamic analysis of the SMAD4-SMAD3-DNA complex at single amino acid resolution. Moreover, the multiplexed uHTS assay demonstrates robustness for screening small molecule inhibitors. Through a pilot screening of an FDA-approved and bioactive compound library, we identified gambogic acid and gambogenic acid as potential hit compounds. These proof-of-concept findings underscore the utility of our optimized multiplexed TR-FRET platform for large-scale screening to discover small molecule inhibitors that target the SMAD4-SMAD3-DNA complex as novel anti-TGFβ signaling agents.

## Introduction

The remarkable clinical achievements of immune checkpoint inhibitor (ICI) therapy have propelled the rapid advancement of immune-mediated anti-tumor strategies, establishing them as the first-line treatment for various tumor types.^1, 2^ Despite the paradigm-shifting progress in cancer immunotherapy over the last decade, the majority of patients fail to respond to current monotherapy based on ICIs, and a significant challenge lies in the occurrence of patient relapse following initial response.^3^ Consequently, there is an urgent and unmet clinical need to address the requirements of the majority of cancer patients, necessitating renewed endeavors to broaden the scope and efficacy of immune system-targeted strategies.

Transforming growth factor-beta (TGFβ) signaling has emerged as a promising target for cancer immunotherapy.^4, 5^ While the role of TGFβ signaling in cancer initiation, progression, and metastasis is multifaceted and context-dependent,^6, 7^ its contribution to the establishment of an immunosuppressive tumor microenvironment (TME) has been extensively documented for both adaptive and innate immune responses.^6, 7^ For instance, TGFβ hampers anti-tumor immunity by inhibiting the proliferation, maturation, differentiation, and activation of natural killer (NK) cells, macrophages, dendritic cells (DCs), and CD8^+^ T cells.^6, 8, 9^ Moreover, it promotes the conversion of naïve CD4^+^ T helper cells into immune suppressive regulatory T (Treg) cells.^10^ Building upon these observations, anti-TGFβ signaling therapies have been actively investigated in clinical trials, particularly in combination with immune checkpoint inhibitors (ICIs), across a wide variety of tumor types.

Numerous therapeutic approaches targeting the anti-TGFβ signaling pathway have been developed, focusing on inhibiting upstream TGFβ and its receptors through the use of neutralizing antibodies, receptor kinase inhibitors, and anti-sense oligonucleotides.^4, 11^ Although promising results have been obtained with these anti-TGFβ therapies in preclinical *in vitro* studies and mouse models, most clinical trials have failed to reproduce these favorable outcomes.^12, 13^ The formidable challenges encountered in the clinical translation of anti-TGFβ signaling therapy not only necessitate the development of novel mechanism-driven strategies but also underscore the imperative to expand the existing repertoire of anti-TGFβ therapeutic options.

SMAD4 (Mothers against decapentaplegic homolog 4) serves as a critical downstream master regulator in the canonical TGFβ signaling pathway.^7^ It functions as a common SMAD (co-SMAD) and acts as an adaptor protein, forming protein-protein interaction (PPI) complex with the receptor-regulated SMADs (R-SMADs), such as SMAD3.^14, 15^ Following the activation of the TGFβ pathway, the SMAD4-SMAD3 PPI complex translocate into the nucleus, binds to SMAD-binding elements (SBE) containing DNA sequences, and initiates the expression of a wide spectrum of TGFβ target genes. However, SMAD4 has been considered ’undruggable’ due to its lack of enzymatic activity and its large protein-protein interaction interface.^16^

To advance the search for SMAD4 inhibitors and broaden the arsenal of anti-TGFβ signaling therapy, we present the development of a multiplexed time-resolved fluorescence resonance energy transfer (TR-FRET) assay capable of simultaneously measuring SMAD4-SMAD3 protein-protein interaction (PPI) and SMAD-SBE protein-DNA interaction (PDI) (Fig. 1). This assay enables the precise monitoring of the dynamics of SMAD4-SMAD3-SBE PPI and PDI at the resolution of single amino acids within a homogeneous cell lysate-based configuration. Furthermore, the assay has been miniaturized and validated for ultrahigh-throughput screening (uHTS) in 1536-well plate format. This optimized and validated uHTS assay will facilitate future large-scale screening campaigns for SMAD4 inhibitors in the quest for novel anti-TGFβ signaling therapy drugs.

**Figure 1.**
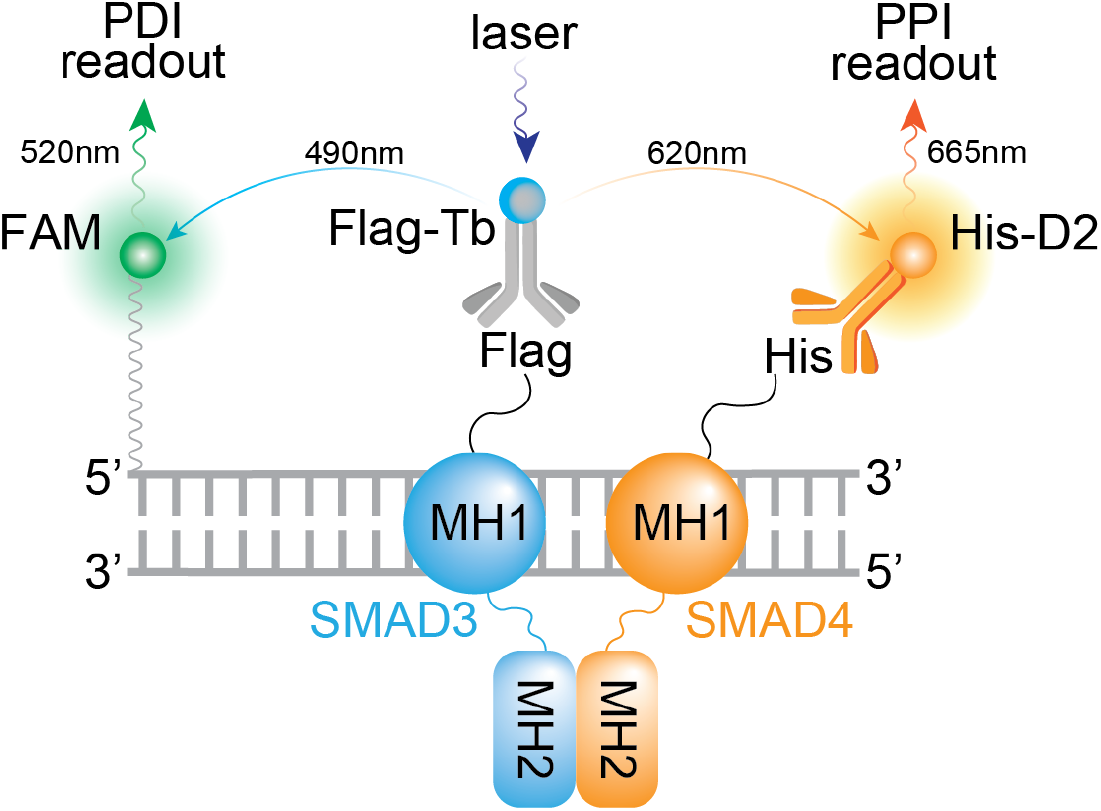
Principle and design of a multiplexed TR-FRET assay. Schematic illustration of the TR-FRET assay with dual-readouts for monitoring SMAD4-SMAD3 protein-protein interaction (PPI) and SMAD3-SBE4 protein-DNA interaction (PDI).

## Results

### Design of the multiplexed SMAD4-SMAD3-DNA TR-FRET assay

In the canonical TGFβ signaling pathway, SMAD4 engages in a cooperative interaction with SMAD3 through its MH2 domain, while its MH1 domain interacts with SBE-containing DNA (Fig. 1).^17^ Thus, in order to expedite the discovery of SMAD4 inhibitors through high-throughput screening (HTS), the development of a robust and scalable bioassay capable of simultaneously monitoring the cooperative dynamics of SMAD4-SMAD3-DNA protein-protein interaction (PPI) and protein-DNA interaction (PDI) is highly desirable. With this objective in mind, we investigated the feasibility of a cell lysate-based time-resolved fluorescence resonance energy transfer (TR-FRET) assay.^18, 19^

TR-FRET is a well-established bioassay widely utilized for monitoring molecular interactions. In essence, TR-FRET signal arises from a proximity-based resonance energy transfer (<10nm) between a long-lived donor fluorophore, such as terbium (Tb), and an acceptor fluorophore with a spectrum that overlaps with the donor. Terbium exhibits a distinctive fluorescence spectrum with four distinct emission peaks at 490, 546, 583, and 620 nm.^20^ This unique fluorescence emission profile of terbium enables the development of multiplexed TR-FRET assays by employing terbium as a single donor paired with multiple spectrally distinct acceptors. This multiplexing capability allows for the simultaneous monitoring of multiple molecular interactions.

To facilitate the monitoring of the SMAD4-SMAD3-DNA complex, we devised a multiplexed TR-FRET assay comprising the following components: 1) Co-expression of His-tagged SMAD4 and Flag-tagged SMAD3 in a cell lysate-based format, 2) Utilization of fluorophore-conjugated anti-fusion-tag antibodies, such as Anti-Flag-Tb and Anti-His-D2, and 3) Employment of synthesized double-stranded DNA (dsDNA) containing SMAD-binding elements (SBEs) that are covalently labeled with a FAM fluorophore (Fig. 1). This assay configuration enables the simultaneous monitoring of SMAD4-SMAD3 PPI and SMAD-DNA PDI through the detection of TR-FRET signals between Tb-D2 and Tb-FAM, respectively.

### Development of the multiplexed SMAD4-SMAD3-DNA TR-FRET assay

To assess the feasibility of the designed assay configuration, we initially optimized the TR-FRET assay using the known SMAD4-SMAD3 PPI as a positive control. According to the assay design, the Tb and D2 fluorophores are brought into close proximity, generating a Tb-D2 TR-FRET signal upon direct interaction between SMAD4 and SMAD3. Consistent with the design, we observed that the cell lysate containing His-tagged SMAD4 and Flag-tagged SMAD3 exhibited significantly higher TR-FRET signals compared to the corresponding negative controls in a concentration-dependent manner (Fig. 2A), with signal-to-background ratios (S/B) reaching up to 30-fold (Fig. 2B). These results not only confirm the feasibility of employing the lysate-based configuration for monitoring the SMAD4-SMAD3 PPI but also establish the optimal lysate concentration for subsequent multiplexing of the SMAD4-SMAD3 PPI with the SMAD-SBE4 protein-DNA interaction (PDI) readout.

**Figure 2.**
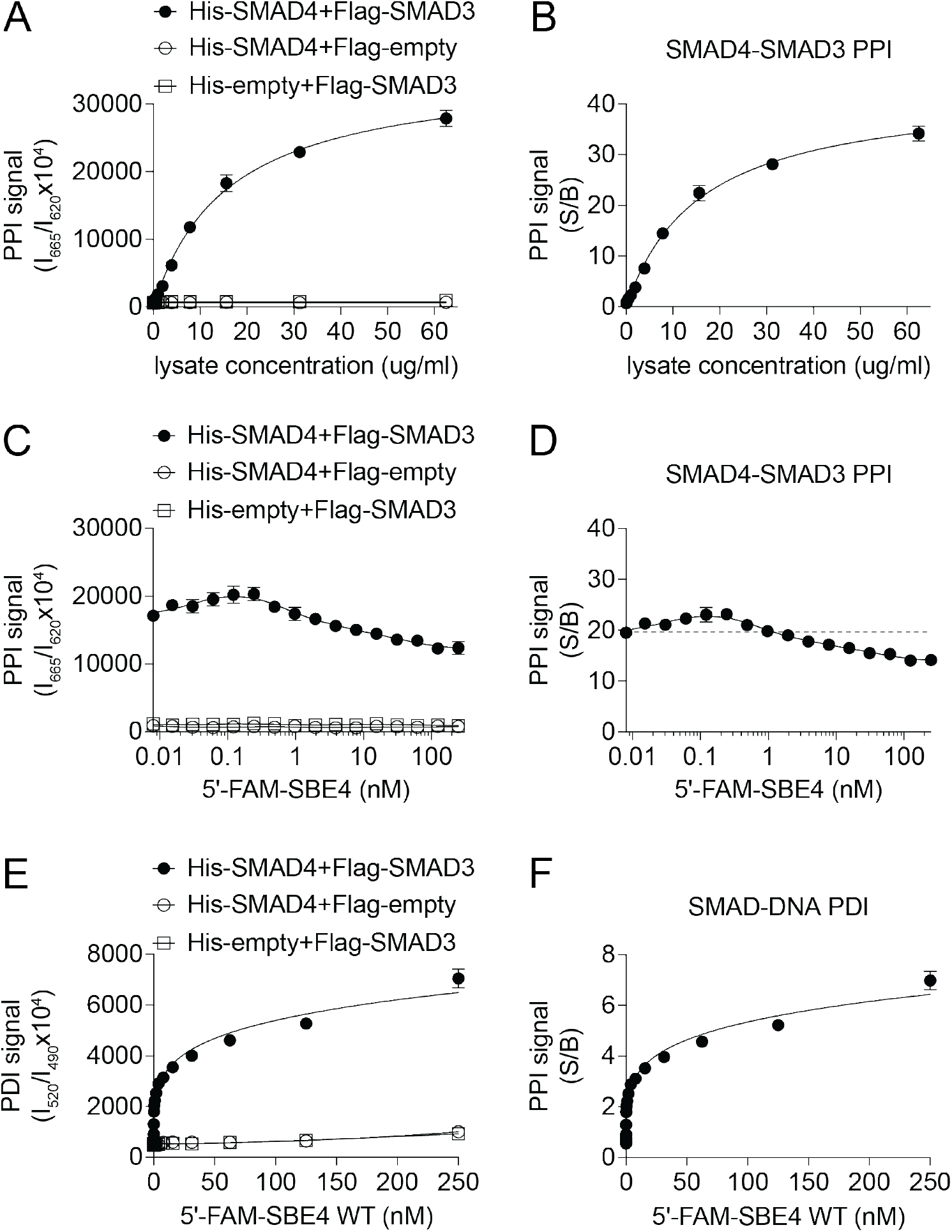
TR-FRET assay development for monitoring the SMAD4-SMAD3-SBE4 complex. **(A-B)** Cell lysate dose-dependent curve of the Tb-D2 TR-FRET PPI signals (A) and S/B from the cell lysate co-expressing His-SMAD4 and Flag-SMAD3 or corresponding controls as indicated. **(C-D)** DNA dose-dependent TR-FRET curve of the Tb-D2 TR-FRET PPI signals (C) and S/B (D) from the lysate co-expressing His-SMAD4 and Flag-SMAD3 or corresponding controls with the titration of 5’-FAM-SBE4 dsDNA as indicated. **(E-F)** DNA dose-dependent TR-FRET curve of the Tb-FAM TR-FRET PDI signals (E) and S/B (F) from the lysate co-expressing His-SMAD4 and Flag-SMAD3 or corresponding controls with the titration of 5’-FAM-SBE4 dsDNA as indicated. The data are expressed as mean ± SD from triplicates from three independent experiments.

To enable the simultaneous monitoring of PPI and PDI, we selected the cell lysate concentration at the EC_60_ (60% maximal effective concentration) condition of the SMAD4-SMAD3 PPI for further evaluation. This concentration was used to test the feasibility of multiplexing the terbium (Tb) fluorophore with other fluorophores that have distinct spectra from D2 (ex/em: 620/665 nm). Initially, we examined a synthesized double-stranded DNA (dsDNA) oligo, 5’-FAM-SBE4, containing four repeats of the SMAD-binding element (SBE) sequence, with 6-carboxyfluorescein (FAM, ex/em: 494/525 nm) conjugated at the 5’-end. As the concentration of 5’-FAM-SBE4 increased, we observed significantly higher PDI signals in the cell lysate containing His-tagged SMAD4 and Flag-tagged SMAD3 compared to the corresponding empty vector controls, in a DNA concentration-dependent manner (S/B > 7-fold) (Fig. 2E-F). Concurrently, from the same samples, we observed stable SMAD4-SMAD3 PPI signals (S/B > 10-fold) (Fig. 2C-D). These results not only demonstrated the feasibility of this multiplexed TR-FRET assay for simultaneous monitoring of SMAD4-SMAD3 PPI and SMAD-DNA PDI from the same sample, but also confirmed the cooperative interactions between SMAD4, SMAD3, and the SBE-containing dsDNA.^17^

### The multiplexed TR-FRET assay is sensitive for detection of SMAD4-SMAD3-DNA complex dynamics

To further assess the assay performance on detecting the dynamic interactions of the SMAD4-SMAD3-DNA complex, we examined the differential PPI and PDI signals using known variants of SMAD4, SMAD3, and DNA that are deficient in complex formation. Initially, we compared the PPI and PDI signals obtained from the cell lysate containing either wild-type (WT) or point mutated (Mut) 5’-FAM-SBE4 double-stranded DNA (dsDNA). As expected, the 5’-FAM-SBE4 Mut dsDNA exhibited a significant reduction of the PDI signal by 45% (p<0.001) and a slight decrease in the PPI signal by 15% (p<0.01) compared to the WT control (Fig. 3A-B).

**Figure 3.**
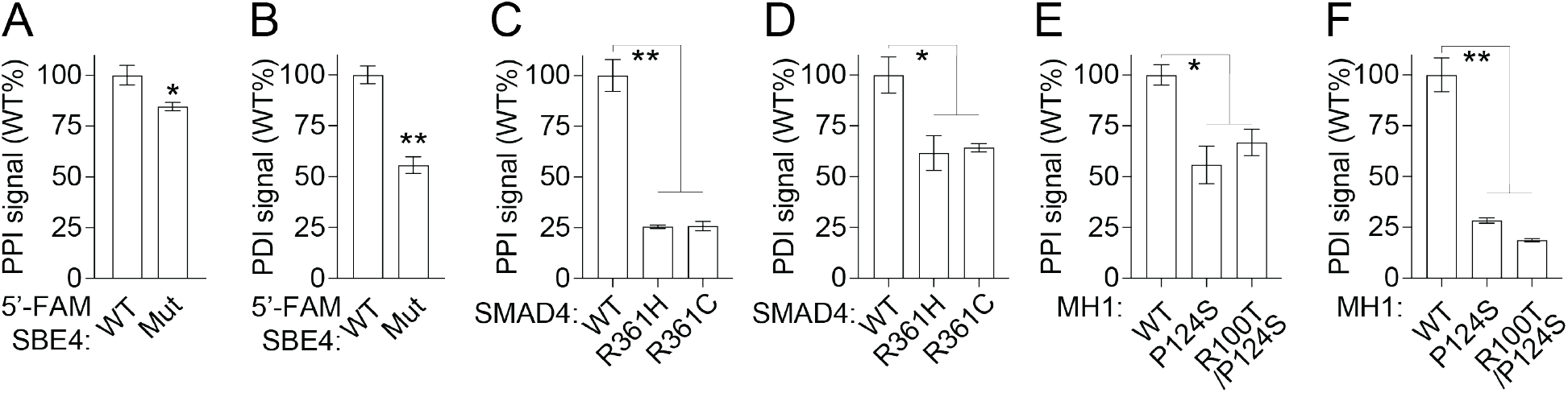
The multiplexed TR-FRET assay allows the detection of SMAD4-SMAD3-DNA complex dynamics. **(A-B)** Bar graphs showing the PPI (A) and PDI (B) signals from cell lysate co-expressing His-SMAD4 and Flag-SMAD3 with 5’-FAM-SBE4 WT or Mut dsDNA. **(C-D)** Bar graphs showing the PPI (C) and PDI (D) signals from cell lysate co-expressing Flag-SMAD3 and His-SMAD4 WT or MH2 domain variants with 5’-FAM-SBE4 WT dsDNA. **(E-F)** Bar graphs showing the PPI (E) and PDI (F) signals from cell lysate expressing SMAD3 and SMAD4 WT or MH1 domain variants with 5’-FAM-SBE4 dsDNA. The data are expressed as mean ± SD from triplicates from three independent experiments. *p<0.01, **p<0.001.

Subsequently, we investigated the PPI and PDI signals using cell lysates expressing SMAD4 R361H or R361C, two naturally occurring cancer-associated hotspot mutations in the MH2 domain that are known to impair SMAD4-SMAD3 PPI.^14, 21, 22^ Our analysis revealed that these SMAD3-binding deficient SMAD4 variants predominantly affected the PPI signal, leading to a substantial reduction of approximately 75% (p<0.001), while exhibiting a slight dampening effect on the PDI signal of approximately 30% (p<0.01) (Fig. 3C-D).

Then, we evaluated the PPI and PDI signals using cell lysates expressing known DNA-binding deficient protein variants with point mutations in the MH1 domain, namely SMAD3 P124S and SMAD4 R100T. Similarly, our findings demonstrated that these DNA-binding deficient variants resulted in a moderate reduction of approximately 25% (p<0.01) in the PPI signal, accompanied by a substantial decrease of approximately 75% (p<0.001) in the PDI signal (Fig. 3E-F).

Collectively, these findings demonstrated the sensitivity of our multiplexed TR-FRET assay, enabling the study of the dynamic formation of the SMAD4-SMAD3-DNA complex at single amino acid/nucleotide variant resolution. It also provided further confirmation that this complex is primarily regulated through the cooperative interaction of the SMAD4-SMAD3 PPI mediated by MH2 domains and the SMAD-DNA PDI facilitated by MH1 domains.^17^

### Assay miniaturization into a 1536-well plate to enable uHTS

To assess the applicability of our multiplexed TR-FRET assay for the discovery of small-molecule modulators, we further miniaturized the assay into a 1536-well plate ultra-high-throughput screening (uHTS) format and evaluated its performance for HTS in terms of signal-to-background ratio (S/B) and Z-prime factor (Z’) (Fig. 4A). The assay demonstrated excellent quality for screening in both 384- and 1536-well plate formats. Robust PPI and PDI signals have been achieved in 1536-well uHTS format with S/B values >15 and ∼6, respectively, along with Z’ values > 0.8 and 0.7 (Fig. 4B-E). These results underscore the feasibility of utilizing our multiplexed TR-FRET assay for conducting uHTS campaigns to identify small molecule modulators targeting the SMAD4-SMAD3-DNA complex.

**Figure 4.**
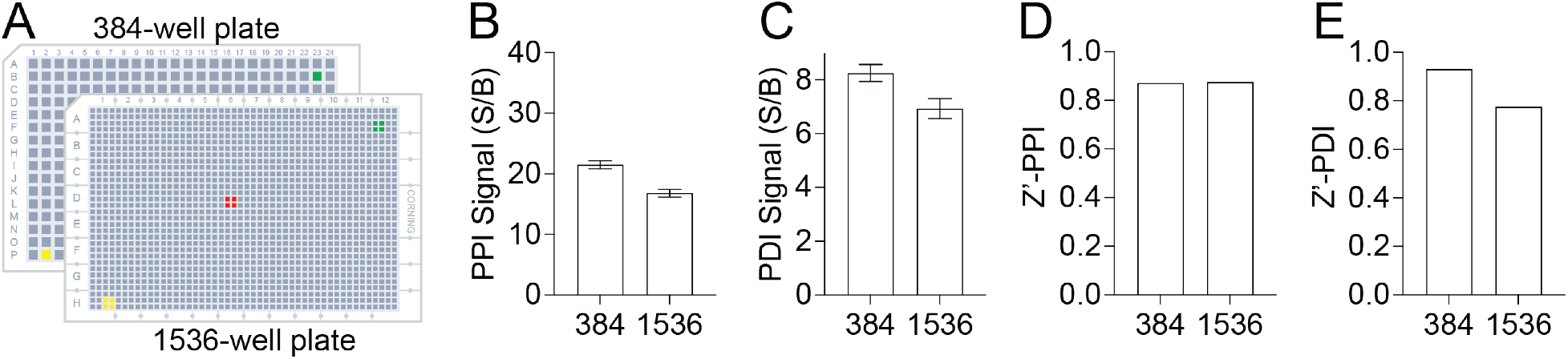
Assay miniaturization for high-throughput screening. **(A)** Schematic illustration of assay miniaturization from 384- to 1536-well plate format. **(B-C)** Bar graphs showing the signal-to-background (S/B) ratio of the PPI (B) and PDI (C) signals for SMAD4-SMAD3-SBE4 WT complex in 384- or 1536-well plate format. **(D-E)** Bar graphs showing the Z prime factor (Z’) of the PPI (D) and PDI (E) signals for SMAD4-SMAD3-SBE4 WT complex in 384- or 1536-well plate format. The data are expressed as mean ± SD from triplicates from three independent experiments.

### Pilot screening for the discovery of small-molecule SMAD4-SMAD3-DNA complex inhibitors

To validate the suitability of our assay for HTS and small-molecule discovery, we conducted a pilot screening using the Emory Enriched Bioactive Library (EEBL), which comprises 12,807 compounds. The primary screening was performed in a 1536-well uHTS format, employing the established conditions described in Figure 4. Each compound was added at a final concentration of 20 μM. For each plate, S/B and Z’ values were calculated for both the PPI and PDI signals. Consistently across ten 1536-well plates, we observed S/B values exceeding 18 and 6, and Z’ values surpassing 0.7 and 0.6 for the PPI and PDI signals, respectively (Fig. 5). These consistent and robust results have validated the excellent performance of our assay in uHTS applications for small-molecule discovery.

**Figure 5.**
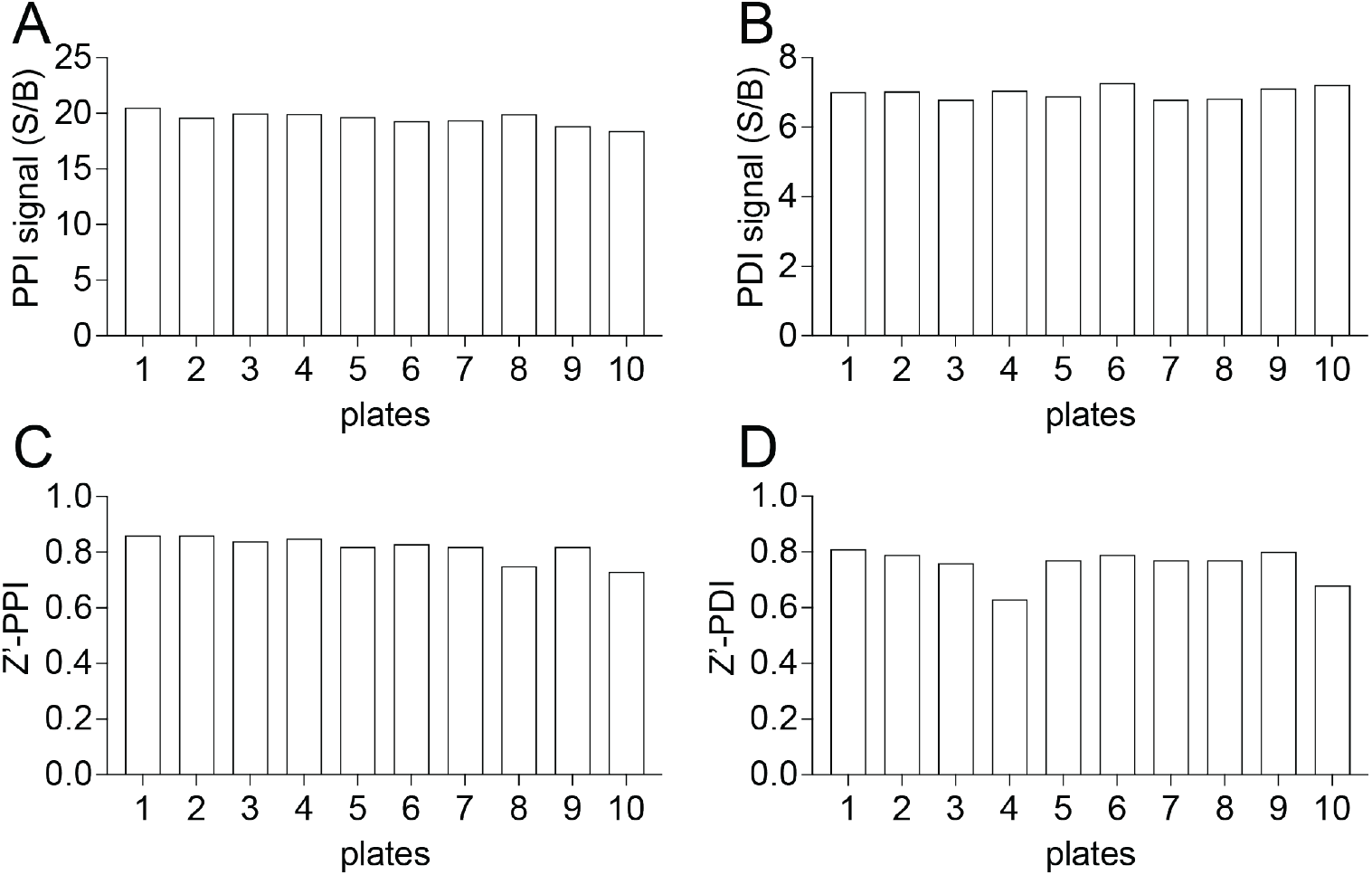
uHTS assay quality control. **(A-B)** Bar graphs showing signal-to-background ratio (S/B) of PPI (A) and PDI (B) readout across 10 plates from the primary screen. The data are expressed as mean calculated from 16 replicates from the primary screen. **(C-D)** Bar graphs showing Z’ of PPI (C) and PDI (D) readout across 10 plates from the primary screen.

The screening results are depicted in Fig. 6A-B. Using a criterion of ≥50% inhibition compared to the DMSO (Dimethyl sulfoxide) control, we identified 251 primary hits from the PPI readout and 212 primary hits from the PDI readout (Fig. 6C). This corresponded to a hit rate of approximately 1.7-2.0%. Considering the cooperative nature of the SMAD4-SMAD3-DNA complex (Fig. 3), we further prioritized 69 hits that showed positive results in both the PPI and PDI readouts (Fig. 6C). Since the TR-FRET assay relies on fluorescence measurements, we additionally prioritized twenty primary hits by excluding assay interference compounds with fluorescence intensity lower or above 20% at the 490 nm, 520 nm, and 620 nm channels, compared to the DMSO control (Fig. 6C).

In the TR-FRET dose-response confirmatory assay using cherry-picked and re-ordered compounds, we confirmed 17 out of the 20 primary hits, which exhibited significant and reproducible effects on decreasing the PPI and PDI signals of the SMAD4-SMAD3-DNA complex (Fig. 6C). These confirmed hits demonstrated potency in modulating the complex formation. The other three primary hits were triaged due to their low potency or lack of consistent effect, indicating they were not suitable candidates for further investigation.

**Figure 6.**
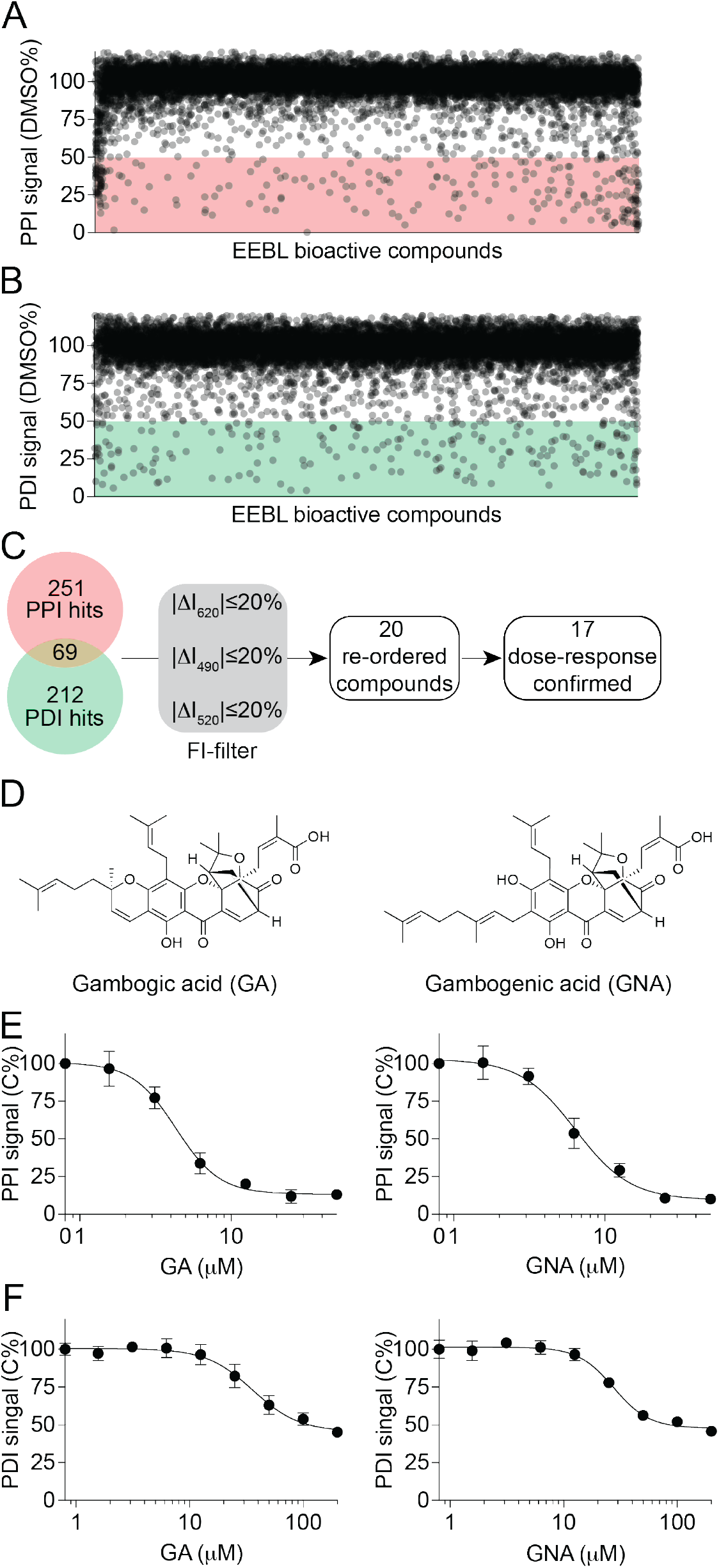
Identification of gambogic and gambogenic acid as SMAD4-SMAD3-SBE4 complex inhibitors from a pilot screening in a 1536-well uHTS format. **(A-B)** Scatter plot showing the PPI (A) and PDI (B) signal induced by compounds from the primary screening. The data are presented as the percentage of the DMSO control from the primary screening. **(C)** Flow chart showing the prioritization of the primary hits. **(D)** Chemical structures of two primary hits, gambogic acid (GA) and gambogenic acid (GNA). **(E-F)** Dose-dependent curves of GA (left) and GNA (right) in inhibiting the PPI (E) and PDI (F) signal. The data are presented as mean ± SD from triplicates from one representative experiments.

### Identification of gambogic and gambogenic acid as SMAD4-SMAD3 PPI inhibitors

Among the 17 hits that were confirmed in the dose-response assay, two structurally similar compounds, gambogic acid (GA) and gambogenic acid (GNA) (Fig. 6D),^23, 24^ attracted particular attention. These two compounds are polyprenylated xanthone natural products derived from the resin of *Garcinia hanburyi*. In the primary screen, both GA and GNA exhibited robust inhibitory effects on the PPI and PDI signals (Fig. 6D). Further dose-response confirmation with re-ordered compounds revealed that GA and GNA induced a concentration-dependent decrease in the PPI signal, with half-maximal inhibitory concentrations (IC_50_) of 4.3 ± 0.5 μM and 6.4 ± 1.6 μM, respectively (Fig. 6E). Moreover, they also induced a concentration-dependent decrease in the PDI signal, with IC_50_ of 36.0 ± 9.4 μM and 27.6 ± 4.8 μM, respectively (Fig. 6F).

It has been previously reported that GA and GNA are cysteine covalent modifiers.^23, 24^ Considering their stronger inhibitory effect on the PPI compared to the PDI in our results (Fig. 6E-F), we formulated a hypothesis that GA and GNA might disrupt the SMAD4-SMAD3-DNA complex by targeting the PPI interface. To investigate this hypothesis, we employed a non-fluorescence affinity-based GST pull-down assay. The results revealed that both GA and GNA dose-dependently decreased the level of SMAD3 protein in the pull-down GST-SMAD4 complex, with IC_50_ values of approximately 11.4 μM and 8.9 μM, respectively (Fig. 7A-B). Furthermore, both GA and GNA significantly inhibited the PPI of SMAD4-SMAD3 MH2 domains, which are the primary domains involved in the PPI, in a dose-dependent manner, with IC_50_ values of approximately 11.4 μM and 9.0 μM, respectively (Fig. 7C-D). These findings suggest that GA and GNA primarily disrupt the SMAD4-SMAD3 PPI by interfering with the interactions occurring at the MH2 domains.

**Figure 7.**
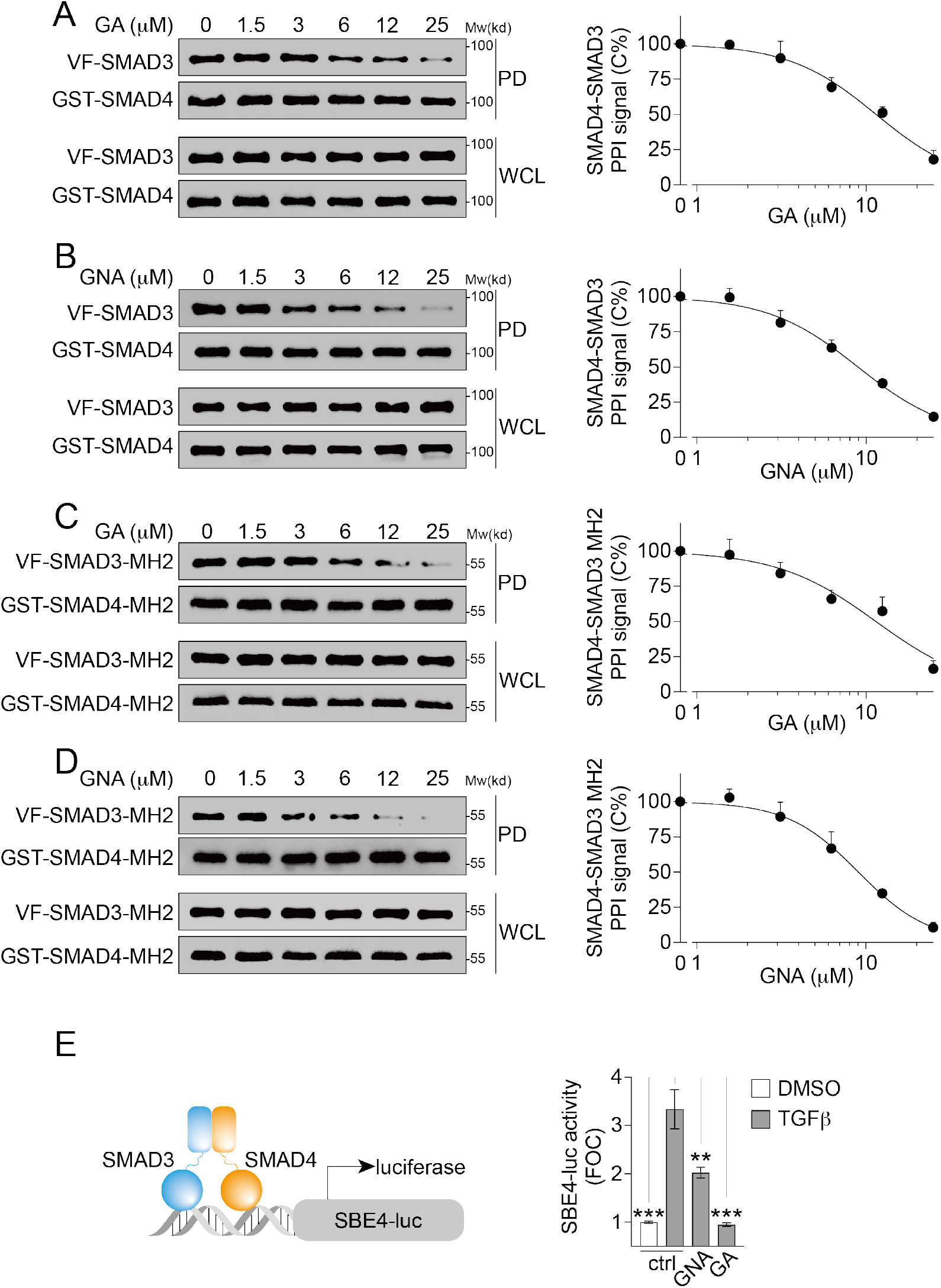
Confirmation of GA and GNA in GST pull-down assay. **(A-B)** Western blot (left) and dose-response curves from the gel quantification (right) showing the inhibition of SMAD4-SMAD3 full-length PPI by GA (A) and GNA (B). The cell lysates expressing the GST-SMAD4 and Venus-flag-tagged (VF) SMAD3 were treated with compound as indicated. Left, the protein samples from the GST pull-down (PD) and the whole-cell lysates (WCL) were analyzed by western blotting. Right, the dose-response curves of the PPI signal were derived from the densitometry analysis of the gels above. The data are presented as mean ± SEM from three independent experiments. **(C-D)** Western blot (left) and dose-response curves from the gel quantification (right) showing the inhibition of SMAD4-SMAD3 MH2 domain PPI by GA (C) and GNA (D). The cell lysates expressing the GST-SMAD4-MH2 and Venus-flag-tagged (VF) SMAD3-MH2 were treated with compound as indicated. Left, the protein samples from the GST pull-down (PD) and the whole-cell lysates (WCL) were analyzed by western blotting. Right, the dose-response curves of the PPI signal were derived from the densitometry analysis of the gels above. The data are presented as mean ± SEM from three independent experiments. **(E)** GA and GNA inhibits TGFβ-induced SBE4-luc reporter activity. HEK293T cells expressing endogenous SMAD4 and SMAD3 were treated with TGFβ (10 ng/mL) and/or GA and GNA for 18 hours as indicated at 5 μM. The TGFβ-induced fold-of-change (FOC) of the luciferase signals are presented as mean ± SD from three independent experiments. **p<0.01, ***p<0.001.

In order to assess the functional impact of GA and GNA on TGFβ signaling, we evaluated the transcriptional activity as a readout for the SMAD4-SMAD3-DNA complex. Specifically, we examined the SBE4-luciferase (luc) activity following TGFβ stimulation (Fig. 7E). Upon TGFβ stimulation, a significant increase in SBE4-luc activity, with a fold-change (FOC) greater than 3, was observed (Fig. 7E). However, treatment with GA or GNA resulted in a significant reduction in TGFβ-induced SBE4-luc activity (Fig. 7E). Taken together, the identification of GA and GNA as positive hits suggests the potential druggability of the SMAD4-SMAD3-DNA complex and paves the way for further exploration in small molecule screening campaigns aimed at discovering novel therapeutics targeting the TGFβ signaling pathway.

## Discussion

DNA-binding transcription factors (TFs) hold immense therapeutic potential in cancer treatment.^25, 26^ However, targeting TFs, other than the nuclear receptor family, poses significant challenges due to their intrinsic disorder and lack of classical binding pockets.^16, 25, 26^ In this study, we have successfully developed a well-designed, optimized, miniaturized and validated multiplexed TR-FRET assay platform to target the SMAD4-SMAD3-DNA complex. This multiplexed assay has demonstrated exceptional sensitivity in investigating the dynamics of the SMAD4-SMAD3-DNA complex at the resolution of single amino acids. It is readily applicable for large-scale small molecule screening campaigns. Through a pilot chemical screen using this assay, we have identified GA and GNA as potential inhibitors of the SMAD4-SMAD3-DNA complex. These compounds not only disrupted the physical protein-protein and protein-DNA interactions within the complex but also effectively inhibited the transcriptional activity induced by TGFβ signaling. These findings from our study underscore the utility of the multiplexed TR-FRET assay for small molecule screening campaigns and provide proof-of-concept evidence supporting the feasibility of directly targeting the previously considered "undruggable" SMAD4-SMAD3-DNA complex. The discovery of small molecule inhibitors for challenging targets like TFs expands the possibilities for therapeutic interventions in cancer and holds great promise for the development of novel anti-cancer treatments.

SMAD4 acts as an adaptor protein, mediating protein-protein interactions (PPI), particularly with R-SMADs like SMAD3, through its MH2 domain.^27^ This interaction enables the translocation of the SMAD4-SMAD3 complex into the nucleus, where the SMADs govern transcription by recognizing DNA sequences containing SBE motifs through their MH1 domain.^28^ The formation of the SMAD4-SMAD3-DNA complex occurs in a highly cooperative manner, playing a crucial role in finely regulated transcriptional processes.^17^ This cooperativity has been extensively documented through low-throughput assays, such as electrophoretic mobility shift assay (EMSA), utilizing purified proteins.^17^ Interestingly, in our cell lysate-based multiplexed TR-FRET assay, we observed that naturally occurring mutations within the MH2 domain of SMAD4 not only disrupt the PPI with SMAD3 (Fig. 3C) but also reciprocally impair the complex formation with DNA (Fig. 3D). Similarly, mutations in the MH1 domain can affect the PPI (Fig. 3E-F). These findings provide further support for the cooperative binding of the SMAD4-SMAD3-DNA complex. Furthermore, our multiplexed TR-FRET platform, as described, holds the potential to be utilized for studying the dynamics of other transcription factor-DNA complexes in a straightforward and quantitative cell lysate-based high-throughput format.

SMAD4 serves as a crucial downstream master regulator of TGFβ signaling, making SMAD4 inhibitors valuable additions to the anti-TGFβ therapy arsenal. However, the development of SMAD4 inhibitors has been challenging due to the protein’s lack of enzymatic activity and its extensive PPI and PDI interfaces. To date, there are no small molecule inhibitors specifically targeting the SMAD4-SMAD3 interaction. Even though an indole derivative, known as SIS3,^29^ has been previously proposed as a potential SMAD3 inhibitor. However, its mode of action involves inhibiting SMAD3 phosphorylation, which indirectly disrupt SMAD4-SMAD3 PPI. In this study, we present a sensitive and robust multiplexed TR-FRET assay capable of simultaneously monitoring the PPI and PDI signals of the SMAD4-SMAD3-DNA complex in cell lysate. This innovative assay enables us to identify small molecule inhibitor candidates, such as GA and GNA, which exhibit the potential to directly interfere with the dynamics of the complex.

Polyprenylated xanthone natural products, such as GA and GNA,^23, 24^ have recently garnered attention due to their diverse biological activities, including anti-cancer, anti-inflammatory, antioxidant, and anti-bacterial effects. However, the polypharmacology properties of these compounds are context-dependent and not yet fully understood.^23, 24^ GA and GNA possess electrophilic characteristics that make them prone to cysteine covalent modification through easy ring opening. Chemoproteomics profiling studies have revealed that GA and its analogs can covalently interact with several protein targets, such as thioredoxins (TRX),^30^ transferrin receptor protein 1 (TFRC),^31^ and serine palmitoyltransferase (SPT),^23^ in a cysteine thiol-dependent manner. In our study, we observed that GA and GNA dose-dependently disrupted the protein-protein interaction of the SMAD4-SMAD3 MH2 domain (Fig. 7C-D). Further investigations are warranted to elucidate whether GA and GNA modulate the SMAD4-SMAD3-DNA complex by interacting with the cysteine residues within the MH2 domain of the proteins.

The identification of polyprenylated xanthones in this study, including GA and GNA, represents the pioneering class of inhibitors targeting the SMAD4-SMAD3 protein-protein interaction. The discovery of these compounds holds great promise as a chemical scaffold for the development of potential clinical candidates, particularly as cancer immunotherapy adjuvants for advanced tumors. The effectiveness of the multiplexed TR-FRET uHTS assay provides the proof-of-concept to support expanded large-scale chemical screening campaigns for discovery novel SMAD4-SMAD3 inhibitors.

## Acknowledgements

We thank members of the Fu/Mo Lab and Emory Chemical Biology Discovery Center for technical support and comments.

## Author Contributions

Conceptualization, X.M., Y.D. and H.F.; Assay Design and Development and chemical screening W.O., Q.N., M.Q., X.M., and Y.D.; Confirmatory assays, W.O. and X.M.; Data Analysis, W.O., Y.D., and X.M.; W.O., and X.M. wrote the initial manuscript; and all were involved in editing.

## Materials and Methods

### Cell Culture

All cell lines were incubated at 37°C in humidified conditions with 5% CO_2_. Human embryonic kidney 293T cells (HEK293T; ATCC, CRL-3216) were maintained in Dulbecco’s Modified Eagle’s Medium (DMEM; Corning, #10-013-CV). Cell culture medium was supplemented with 10% fetal bovine serum (ATLANTA biologicals, #S11550) and 100 units/mL of penicillin/streptomycin (Cell Gro, Cat# 30-002-CI).

### Molecular cloning and mutagenesis

The WT SMAD3 (Clone# IOH27044) and SMAD4 (Clone# IOH3638) genes in pDONR221 plasmid were gifted from Drs. Gordon Mill and Yiu Huen Tsang at Oregon Health Science University. The SMAD4 point mutations, including R361H, R361C and R100T, along with SMAD3 point mutation P124S, were introduced using QuikChange Lightning Site-Directed Mutagenesis Kit (Agilent Technologies) and respectively the SMAD4 or SMAD3 pDONR221 plasmid as DNA template and the following primers: R361H forward primer (5’-CTTCTGGAGGAGATCACTTTTGTTTGGGTCAAC-3’), R361C forward primer (5’-CCTTCTGGAGGAGATTGCTTTTGTTTGGGTCAA-3’), R100T forward primer (5’-TGCCCGTCTCTGGACGTGGCCTGATCTTCA-3’), P124S forward primer (5’-AGGTCTGCGTGAATTCCTACCACTACCAGA-3’), and corresponding reverse complementary primers. The SMAD4 and SMAD3 MH2 domain truncation plasmids were generated by PCR using Platinum™ *Taq* DNA Polymerase kit (Thermo Fisher Scientific) and respectively the SMAD4 or SMAD3 pDONR221 plasmid as DNA template and the following primers: SMAD4 MH2 forward primer (5’-GGGGACAAGTTTGTACAAAAAAGCAGGCTTCGAAGGAGATAGAACCATGGATATGGCTCCTGAGTATTGGTGTTCCATT-3’), SMAD4 MH2 reverse primer (5’-GGGGACCACTTTGTACAAGAAAGCTGGGTTGTCTAAAGGTTGTGGGTCTGCAAT-3’), SMAD3 MH2 forward primer (5’-GGGGACAAGTTTGTACAAAAAAGCAGGCTTCGAAGGAGATAGAACCATGGATATGTTGGACCTGCAGCCAGTTACC −3’), SMAD3 MH2 reverse primer (5’-GGGGACCACTTTGTACAAGAAAGCTGGGTTAGACACACTGGAACAGCGG −3’). Gateway cloning (Invitrogen) was used to generate GST-tagged, Venus-flag-tagged, Flag-tagged and 6XHis-tagged plasmids as previously described.^14, 18, 19, 32–34^ The vector backbones are pDEST27 vector (Invitrogen) for GST-tag, pDEST26 (Invitrogen) for 6XHis tag, pSCM167 for Venus-flag-tag, pcDNA3.2-V5-dest for Flag-tag constructs. All plasmids generated were confirmed by sequencing.

### Multiplexed Time-resolved fluorescence resonance energy transfer (TR-FRET) assay

TR-FRET assays were performed using cell lysate from HEK293T cells expressing Flag-tagged SMAD3 and His-tagged SMAD4 WT or MUT proteins. The FRET buffer used throughout the assay contains 20 mM Tris-HCl, pH 7.0, 50 mM NaCl, and 0.01% nonidet P-40 (NP-40). Briefly, the HEK293T cells were transiently co-transfected with Flag-tagged SMAD3 WT (1.5 μg/well) or MUT (1.5 μg/well) and His-tagged SMAD4 WT (1.5 μg/well) or MUT (1.5 μg/well) plasmids in the 6-well plate. FuGENE (FuGENE, Promega, Cat# E2920) were used as transfection reagent at 3:1 (FuGENE/plasmid mass) ratio. Forty-eight hours after transfection, cell lysates were prepared in 200 μl 1% NP-40 lysis buffer containing 150 mM NaCl, 10 mM HEPES pH 7.5, 1% nonident P-40 (IGEPAL CA-630, Sigma-Aldrich), 5 mM sodium pyrophosphate, 5 mM NaF, 2 mM sodium orthovanadate, 10 mg/L aprotinin, 10 mg/L leupeptin and 1mM PMSF.

To determine the optimal cell lysate concentration for HTS, the cell lysate concentration dependent TR-FRET assay was performed in black 384-well plates (Corning Costar, #3573). 15 μL of stock cell lysate was 2-fold serially diluted in FRET buffer and mixed with 15 μL mixture of fluorophore-conjugated antibodies. The total volume for each well was 30 μL containing the cell lysate, anti-FLAG M2-Tb cryptate antibody (Cisbio 61FG2TLF, 1:1000 dilution) and anti-6XHIS-D2 antibody (Cisbio 61HISDLF, 1:500 dilution). The plate was centrifuged at 500 g for 5 min and incubated at 4 °C for 30 min. TR-FRET signals for PPI (protein-protein interaction) were measured using the BMG Labtech PHERAstar FSX reader with the HTRF optic module (excitation at 337 nm, emission A at 665 nm, emission B at 620 nm, integration start at 50 μs, integration time for 150 μs and 8 flashes per well). All FRET signals were expressed as a TR-FRET ratio: F665nm /F620nm x 10000.

To determine the optimal fluorescein-tagged oligo concentration for HTS, the oligo concentration dependent TR-FRET assay was performed in black 384-well plates (Corning Costar, #3573). 15 μL of stock oligo mixture containing pre-diluted cell lysate (with the desired concentration) and fluorescein-tagged oligos (500 nM) was 2-fold serially diluted in the same concentration pre-diluted cell lysates, and mixed with 15 μL mixture of fluorophore-conjugated antibodies. The total volume for each well was 30 μL containing the fluorescein-tagged oligos, cell lysate, anti-FLAG M2-Tb cryptate antibody (Cisbio 61FG2TLF, 1:1000 dilution) and anti-6XHIS-D2 antibody (Cisbio 61HISDLF, 1:500 dilution). The plate was centrifuged at 500 g for 5 min and incubated at 4 °C for 30 min. TR-FRET signals for PDI (protein-DNA interaction) were measured using the BMG Labtech PHERAstar FSX reader with the HTRF optic module (excitation at 337 nm, emission A at 520 nm, emission B at 490 nm, integration start at 50 μs, integration time for 150 μs and 8 flashes per well). TR-FRET signals for PPI were measured using the BMG Labtech PHERAstar FSX reader with the HTRF optic module (excitation at 337 nm, emission A at 665 nm, emission B at 620 nm, integration start at 50 μs, integration time for 150 μs and 8 flashes per well). PDI FRET signals were expressed as a TR-FRET ratio: F520nm /F490nm x 10000. PPI FRET signals were expressed as a TR-FRET ratio: F665nm /F620nm x 10000.

### uHTS TR-FRET screening for small molecule PPI inhibitor discovery

Ultra-high-throughput screening (uHTS) for small molecule PPI inhibitor discovery was performed using the TR-FRET assay in the black 1536-well plate (Corning Costar, #3724) with a total volume of 5 μL in each well. The amount of cell lysate and antibodies were scaled down proportionally from the conditions with the optimal assay window identified from 384-well plate. Briefly, 5 μL solutions containing cell lysate and antibodies at desired concentrations were dispensed in the 1536-well plate using a Multi-drop Combi Reagent Dispenser (ThermoScientific). The last column was used as the empty vector background control. Subsequently, the Emory Enriched Bioactive Library (EEBL) compounds (100 nL) were added into wells in each plate using Biomek NXP Automated Workstation (Beckman) from a compound stock plate to give the final concentration of 20 μM. The final DMSO concentration was 2% (v/v) in samples with compound treatment. Each sample was tested with single point. After overnight incubation at 4 °C, FRET signal was measured using the BMG Labtech PHERAstar FSX reader with the HTRF optic module. To evaluate the performance of the assay for HTS, Z’ factor and signal-to-background(S/B) ratio were calculated for the TR-FRET titration experiment according to the following equations:

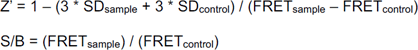

where SD_sample_ and SD_control_ are standard deviations, and FRET_sample_ and FRET_control_ represent the TR-FRET signal (PPI or PDI, respectively) from lysate sample containing oligos, His-SMAD4 and Flag-SMAD3, or oligos and empty Flag-vector controls, respectively. S/B suggests the signal window of the assay and Z’ factor reflects the robustness of the assay for HTS. A Z’ factor between 0.5 and 1 indicates a robust assay and suit for HTS. Screening data were analyzed using Bioassay software from CambridgeSoft (Cambridge, MA). The effect of compound on PPI and PDI modulation was quantified as the change of TR-FRET signal (ΔTR-FRET) upon compound treatment using the equation 100 * (FRET_compound_ - FRET_DMSO_)/FRET_DMSO_, where FRET_compound_ and FRET_DMSO_ are the TR-FRET signals from PPI or PDI in the presence of library compound or DMSO with background FRET_vector_ subtracted. Cutoff of ΔTR-FRET ≥ 50% was used to prioritize the positive hits.

### GST pull-down assay

To validate the hits from the pilot screening, we performed orthogonal GST pull-down assay using cell lysate from the HEK293T cells transfected with Venus-flag-SMAD3 and GST-SMAD4, or Venus-flag-SMAD3-MH2 and GST-SMAD4-MH2. After 48hrs of transfection, the cells were lysed in the 1% NP-40 lysis buffer and incubated with compounds for 16 hours rotating at 4°C, and then incubated with glutathione-conjugated beads (GE 17527901) for 2h at 4°C. Beads were washed twice with the 1% NP-40 lysis buffer and eluted by boiling in sodium dodecyl sulfate (SDS) sample buffer (Thermo Fisher Scientific) and subjected to western blot analysis.

### Western blot

Proteins in the SDS sample buffer were resolved by 10% SDS polyacrylamide gel electrophoresis (SDS-PAGE) and were transferred to nitrocellulose filter membranes at 100 V for 2h at 4°C. After blocking the membranes in 5% nonfat dry milk in TBST (20mM Tris-base, 150mM NaCl, and 0.05% Tween 20) for 1 hour at room temperature, membranes were blotted with the indicated antibodies at 4°C overnight. Membranes were washed by 1xTBST for three times, 15 minutes each time. SuperSignal West Pico PLUS Chemiluminescent Substrate (Thermo, #34580) and Dura Extended Duration Substrate (Thermo, #34076) were used for developing membranes. The luminescence images were captured using ChemiDoc^TM^ Touch Imaging System (Bio-Rad).

### SMAD binding element (SBE) luciferase reporter assay

The SMAD3/SMAD4 complex transcriptional activity was measured using the SBE-luciferase reporter system. HEK293T were used to measure the luciferase activity. Cells were plated in 6-well plate and co-transfected with SBE4-Luc plasmid (1 μg, Addgene,16495) and pDEST26-Renilla plasmid (0.1 μg) using FuGENE® HD (Promega, Cat# E2312). Twenty-four hours after transfection, the cells were pre-treated with gambogic acid or gambogenic acid at 5 μM was 6h followed by TGFβ (10 ng/mL) stimulation for additional 18h. Renilla and Firefly luciferase activities were measured by Envision Multilabel plate reader (PerkinElmer) using a Dual-Glo luciferase kit (Promega, Cat# E2920) according to the manufacturer’s instructions. The normalized luminescence was calculated as the ratio of luminescence of Firefly luciferase over the luminescence of Renilla luciferase.

